# Microcrystalline hydroxyapatite is not inferior to fluorides in clinical caries prevention: a randomized, double-blind, non-inferiority trial

**DOI:** 10.1101/306423

**Authors:** Ulrich Schlagenhauf, Karl-Heinz Kunzelmann, Christian Hannig, Theodor W. May, Helmut Hösl, Mario Gratza, Gabriele Viergutz, Marco Nazet, Sebastian Schamberger, Peter Proff

## Abstract

Recent evidence for a significant interference of microcrystalline hydroxapatite (HAP) particles with re- and demineralisation processes at the tooth-biofilm interface suggested, that they may be promising candidates for efficacious caries prevention. This multicenter randomized controlled non-inferiority trial evaluated the impact of the 2 x daily use of a HAP dentifrice without fluoride on the progression of enamel caries in adolescent caries-risk patients subjected to orthodontic therapy, with a fluoridated AmF/SnF dentifrice serving as a positive control. Primary study outcome was the occurrence of enamel caries lesions ≥ ICDAS (*International Caries Detection and Assessment System*) code 1 around orthodontic brackets on the vestibular surfaces of teeth 15-25 within the 168 days observation period. Secondary study outcomes were the occurrence of enamel caries lesion ≥ ICDAS code 2, Plaque Index (PlI) and Gingival Index (GI). Out of 150 recruited patients, 147 were included in the intent to treat analysis (ITT); 133 finished the study per protocol (PP). PP data analysis revealed the occurrence of enamel caries ≥ ICDAS code 1 in 54.7% of the HAP group patients compared to 60.9% of the fluoride control. In the ITT analysis the corresponding numbers were 56.8% (HAP) and 61.6% (control). Non-inferiority testing of the ITT as well as the PP data set proved that the caries preventive efficacy of the HAP dentifrice was not inferior to the protection provided by the fluoridated AmF/SnF control. Regarding all assessed secondary outcomes (enamel caries ≥ ICDAS code 2, GI, PlI) no significant differences between both experimental groups were observed. Within the restraints set by design and study population of this trial microcrystalline HAP as ingredient of toothpaste may thus be regarded a promising supplement to fluorides in clinical caries prevention (ClinicalTrials.gov: NCT02705456).

## Introduction

In recent years findings, mostly derived from *in vitro* studies, suggested, that microcrystalline hydroxyapatite (HAP) particles may be promising candidates for the prevention of cariogenic demineralization and the stimulation of remineralization processes on enamel and dentine surfaces [1–3]. Huang et al. (2011) reported a regain of mineral content and an increase in microhardness on demineralized bovine enamel slabs that had subsequently been exposed to microcrystalline HAP particles [2]. The observed increase in mineral content proved to be pH-dependent and was significantly higher under acidic conditions. Lin et al. (2014) discovered a significant inhibition of future demineralisation under acidic conditions after HAP application due to the formation of a protective HAP layer over the prism-prism sheath interfaces, where enamel dissolution usually is initiated [4]. In an *in situ* - study the use of a zinc-carbonate HAP microcluster-containing mouth rinse significantly reduced bacterial colonization on bovine enamel slabs worn intraorally by healthy volunteers [5].

Hannig and Hannig (2010) put these *in situ* and *in vitro* findings into a more comprehensive perspective by stating that already physiological tooth wear constantly releases HAP particles into the oral environment, which may subsequently interfere with de- and remineralisation processes as well as with the metabolism of the oral microbiota at the tooth-bacterial biofilm interface [6]. The impact of microcrystalline HAP as an ingredient of dentifrices has been positively evaluated in controlled clinical trials regarding dentinal hypersensitivity [7–10] and parameters of periodontal health [11]. Up to date, however, comparable data regarding the caries-preventive properties of HAP toothpastes are mostly missing. They are limited to positive findings from *in situ* studies on extracted teeth or standardized enamel and dentine specimen, being subjected to different toothpaste slurries and worn in between by volunteers in intraoral appliances [12–15]. As orthodontic therapy with fixed appliances is known to be associated with an increased incidence of the overgrowth of a caries-promoting microbiota [16] and the development of white spot enamel caries lesions [17–19], this study aimed at the assessment of the caries-preventive impact of the regular use of a fluoride-free HAP dentifrice in this particular group of caries risk patients. Due to the abundant evidence for the caries preventive efficacy of fluorides [20, 21], clinical caries studies may no longer involve a true negative control for obvious ethical reasons. Thus a non-inferiority trial was conducted. The study hypothesis to be tested was, that, in terms of caries prevention, the regular use of the HAP test dentifrice is not inferior to the regular use of a fluoridated control with proven efficacy.

## Material and methods

The investigation was designed as a multicenter, prospective, parallel group, two arm, double-blind, randomized clinical non-inferiority trial to be performed at the German study centers Wuerzburg (leading study center), Regensburg, Munich, Dresden and Frankfurt. The study protocol was prepared in accordance with the declaration of Helsinki and met the criteria of GCP. It was approved by the ethics committee of the University of Wuerzburg (file #184/13) on March 28^th^, 2013. Registering clinical trials was not yet generally regarded a mandatory prerequisit to be performed prior to study initiation in 2012 and 2013 during the planning phase of this investigation. Therefore, registration with ClinicalTrials.gov (identifier: NCT02705456) was performed late on February 25^th^, 2016 in the final phase of patient recruitment, which had started already more than 1 year earlier. The authors confirm that all ongoing and related trials for this drug/intervention are registered.

### Study design

The design of the study is schematically depicted in Fig 1.

**Fig 1.**
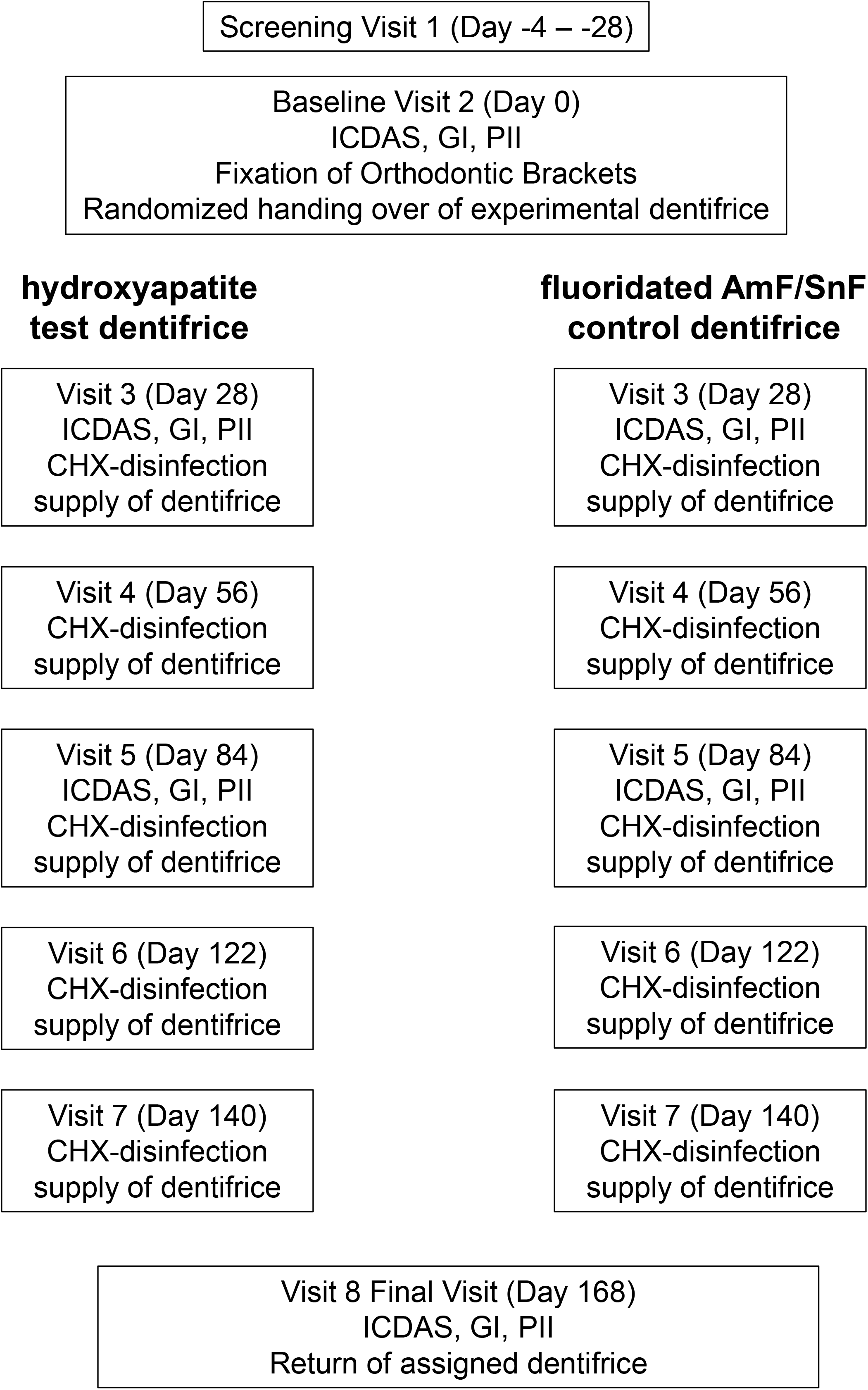
Study Design. At visit 1 (-4 to -28 days prior to baseline) patients scheduled for orthodontic therapy were screened for study eligibility. Those meeting it and giving informed consent were subsequently scheduled for the baseline visit 2 (day 0). At visit 2 Plaque Index (PlI) as well as Gingival Index (GI) scores were recorded from the vestibular surfaces of teeth 15 to 25 followed by professional tooth cleaning and the subsequent assessment of the vestibular enamel surfaces of teeth 15 to 25 according to ICDAS II criteria. Afterwards orthodontic brackets were adhesively mounted to the vestibular surfaces and any excess of adhesive resin was removed. No sealants, fluoride varnishes or any other caries-preventive layers surrounding the brackets were applied. Using a randomization list a supply of either the test dentifrice or the control dentifrice, calculated to be adequate for 4 weeks of 2 x daily repeated toothbrushing, as well as a standardized electric tooth brush (Oral-B Pulsar 35; Procter & Gamble GmbH, Germany) to be used for the duration of the study, were handed over to the study patients. The dosage of the assigned dentifrice (2 x daily a streak of approx. 1 g) and the use of the electric tooth brush were practically instructed and the patients informed, to bring back all assigned toothpaste tubes at the next scheduled visit. At day 28 the sequence of recording PlI, GI and ICDAS II scores was repeated as described for visit 2. As an additional caries-preventive measure, teeth 15-25 were disinfected with a topically applied 1% chlorhexidine (CHX) gel. Toothpaste tubes supplied at visit 2 were taken back and a new supply for the next 4 weeks handed over. At day 56 (visit 4) oral hygiene was reinstructed and the cleaning/disinfection procedures as well as the return/handing over of the toothpaste supply performed as described before. At day 84 (visit 5) the recording of GI, PlI and ICDAS II scores as well as cleaning and disinfection were repeated as described before. Next to a new supply of toothpaste also a new Pulsar 35 electric toothbrush was handed over. At day 112 (visit 6) and at day 140 (visit 7) procedures performed were identical to those at day 56 (visit 4). At day 168 (visit 8) the final assessment of PlI, GI and ICDAS II scores as well as the return of the study dentifrices was conducted as described before. Furthermore, at each study visit patients were asked about the occurrence of important harms or unintended effects related or unrelated to the use of the study dentifrices.

### Study population

The trial was performed in adolescents and young adults being scheduled for orthodontic therapy with fixed appliances.

#### Inclusion criteria were

- age 11-25 yrs.
- scheduled orthodontic therapy with fixed appliances of at least 6 months duration comprising the placement of orthodontic brackets on the vestibular surfaces of teeth 15 - 25
- regular (2x daily) oral home care with toothbrush and toothpaste
- caries promoting salivary counts of mutans streptococci ≥ 10^5^ CFU/ml, which were determined using the CRT^®^ bacteria test (Ivoclar Vivadent, Liechtenstein) [22].

#### Exclusion criteria were

- untreated caries lesions of ICDAS code 3-6 on any tooth
- treated carious lesions of ICDAS code 3-6 on the vestibular surfaces of teeth 15-25
- diseases or conditions interfering with the salivary flow or requiring the regular use of medications interfering with it
- antibiotic therapy within the last 6 weeks before study participation or necessity for antibiotic prophylaxis during dental interventions
- known allergies to ingredients of the experimental dentifrices

### Interventions - experimental dentifrices

#### a) Test Dentifrice

The test dentifrice (Karex^®^ Zahnpasta; Dr. Kurt Wolff GmbH & Co. KG, Germany) was provided by the sponsor of the study. It contained 10% of microcrystalline HAP as the main caries-preventive agent and also the following ingredients: Aqua, glycerol, hydrogenated starch hydrolysate, xylitol, hydrated silica, Silica, aroma, cellulose gum, sodium methyl cocoyl taurate, *Helianthus anuus* seed oil, polyglyceryl- 3 palmitate, polyglyceryl-6 caprylate, *Usnea barbata* extract.

#### b) Control Dentifrice

A commercially available fluoridated toothpaste (meridol^®^ Zahnpasta; CP GABA GmbH, Germany) was used as a positive control. It contained amine fluoride and stannous fluoride in concentrations of 350 ppm and 1050 ppm, respectively, and furthermore the following ingredients: Aqua, sorbitol, hydrated silica, silica dimethyl silylate, hydroxyethylcellulose, PEG-40, hydrogenated castor oil, cocamidopropyl betaine, aroma, sodium gluconate, PEG-3 tallow aminopropylamine, saccharin, hydrochloric acid, potassium hydroxide, CI 74160.

## Primary outcome

The primary outcome was set to the percentage of subjects in each experimental group exhibiting the new occurrence of at least one enamel caries lesion of ICDAS code 2 or higher on any vestibular surface of the 10 evaluated teeth 15 to 25 during the observation period of 168 days.

## Assessment of carious lesions

The occurence of caries lesions was evaluated visually on the vestibular surfaces of teeth 15 to 25 according to the criteria of the *International Caries Detection and Assessment System* (ICDAS-II) [23]. The examination was performed at baseline, prior to the fixation of the orthodontic brackets, and was repeated 28 days, 84 days and 168 days later. All teeth were professionally cleaned before each assessment from any adhering bacterial biofilms or stains. The development of a caries lesion > ICDAS code 3 during the course of the study on any tooth and observed at any visit was defined as an immediate study exit criterion.

### Interexaminer reliability

To ensure interexaminer reliability, prior to the study onset all examiners were instructed to pass the ICDAS e-learning course at the icdas.org website and were subsequently trained in person by an experienced expert (K.H.K.) to perform ICDAS assessments in reference patients. Grading skills were retrained 3 times during the course of the study using another internet-based ICDAS training tool. Similar to Luz et al. [24] it confronted the examiners with a random sample of 40 pictures of upper premolars, canines and incisors with surface integrities ICDAS codes 0-3. 50% of the pictures of a given sample were randomly presented in duplicates to evaluate the ability of the examiners to reproduce their own assessments.

Interrater reliability analysis revealed a mean weighted kappa = 0.75 for the first assessment run, which increased to kappa = 0.80 for the final calibration, indicating “substantial agreement” among the different examiners throughout the study [25]. Although up to three examiners were trained and calibrated at each study center before the onset of the trial, at four centers the bulk of the practical evaluations was performed by a single principal examiner (Munich 100% of all visits, Frankfurt 100%, Regensburg 96 %, Wuerzburg 96 %) At the center in Dresden the principal examiner performed 58% of all examinations, the second examiner 38%.

### Secondary outcomes

Secondary outcomes were plaque coverage and gingival inflammation at baseline and at day 168 evaluated by recording the

- Plaque Index (PlI) [26] and the
- Gingival Index (GI) [27], respectively.

## Sample size calculation

Based on a reported caries incidence rate of about 60% in a preceding caries trial assessing orthodontic patients with fixed braces, who were not beeing preselected for particular caries-promoting risk factors [18], the likelihood for the occurrence of an ICDAS code 2 lesion during the 168 day observation period in this cohort of caries-risk individuals with elevated salivary numbers of caries-promoting mutans streptococci was extrapolated to be p=80% for the control group using the fluoridated toothpaste. The difference between both experimental groups not be regarded clinically relevant was set to Δ ≤ 20%. A sample size of 2 x 74 study patients was calculated to be sufficient to reject the null hypothesis, that the test dentifrice is inferior to the control dentifrice, using a non-inferiority margin of Δ = 20% for the primary outcome measure and one-sided, exact Fisher Test (α = 5%, power = 80%).

## Blinding, randomisation

The trial was designed to blind study patients and examiners to the group assignment. For this purpose, both study dentifrices (test/control) were filled into neutral plastic tubes of identical shape and color by an independent, GMP certified laboratory for cosmetics. Using block randomization with a block size of 4 a random list was generated to code-label test and control tubes with consecutive unique identification numbers. Randomization of dentifrice assignment was stratified by study center. Handing out of the experimental dentifrices to the study patients followed the sequence of the identification numbers and was performed by trained study nurses not involved in the examination of the study participants. To maintain blinding of examiners and study patients, the study patients were instructed not to discuss toothpaste-related issues with the examiners but with the study nurses only, who were also responsible for instructing the patients in efficacious oral hygiene and taking back the empty or unused dentifrice tubes at the subsequent visits. The number of study nurses varied between a minimum of one and a maximum of four per study center.

## Statistical analysis

The primary outcome measure was analysed primarily for the PP population and repeated for sensitivity reasons, for the ITT population. The exact confidence limits (Clopper-Pearson) were computed to test non-inferiority (cp. [28]). For the primary outcome measure, non-inferiority was claimed, if the upper limit of the one-sided 95% confidence for the corresponding difference between test and control dentifrice was less than Δ ≤ 20%.

In addition, two-sided Wilcoxon-Mann-Whitney tests were used for between group comparisons and Friedman tests for within group comparisons for secondary outcomes.

SAS^®^ 9.3 software package (SAS Institute Inc., USA) was used for statistical evaluations.

## Patient Recruitment

Out of a total of 281 screened individuals, 150 patients meeting the inclusion criteria gave written informed consent and were recruited at the study centers in Wuerzburg (n=36), Regensburg (n=72), Dresden (n=28), Munich (n=12) and Frankfurt (n=2). The first patient was included in the trial on November 13^th^, 2013, the last patient left the trial on August 28^th^, 2016. At the study centers Wuerzburg, Regensburg and Dresden not only center patients but also orthodontic patients being treated in private practice were included and assessed by the examiners of the center.

## Dropouts

Six patients of the test group and 4 patients of the control group terminated study participation prematurely due to lack of interest or not keeping the follow-up appointments. Further 6 patients completed the study but were excluded from the PP analysis due to insufficient dosing of the assigned dentifrice, calculated from the residual weight of the returned dentifrice tubes. All but one patient of the test group and all patients of the control group received at least one dose of the assigned dentifrice (n=149) and were thus primarily included in the ITT analysis set. As two study patients left the trial already before the first reevaluation at week 4, the total number of study individuals suitable for an inclusion in the ITT analysis of caries development further decreased to n=147. No important harms or unintended effects related or unrelated to the use of the study dentifrices were reported. Finally, a total of 133 study patients (64 test / 69 control) was included in the PP analysis set (Fig 2).

**Fig 2.**
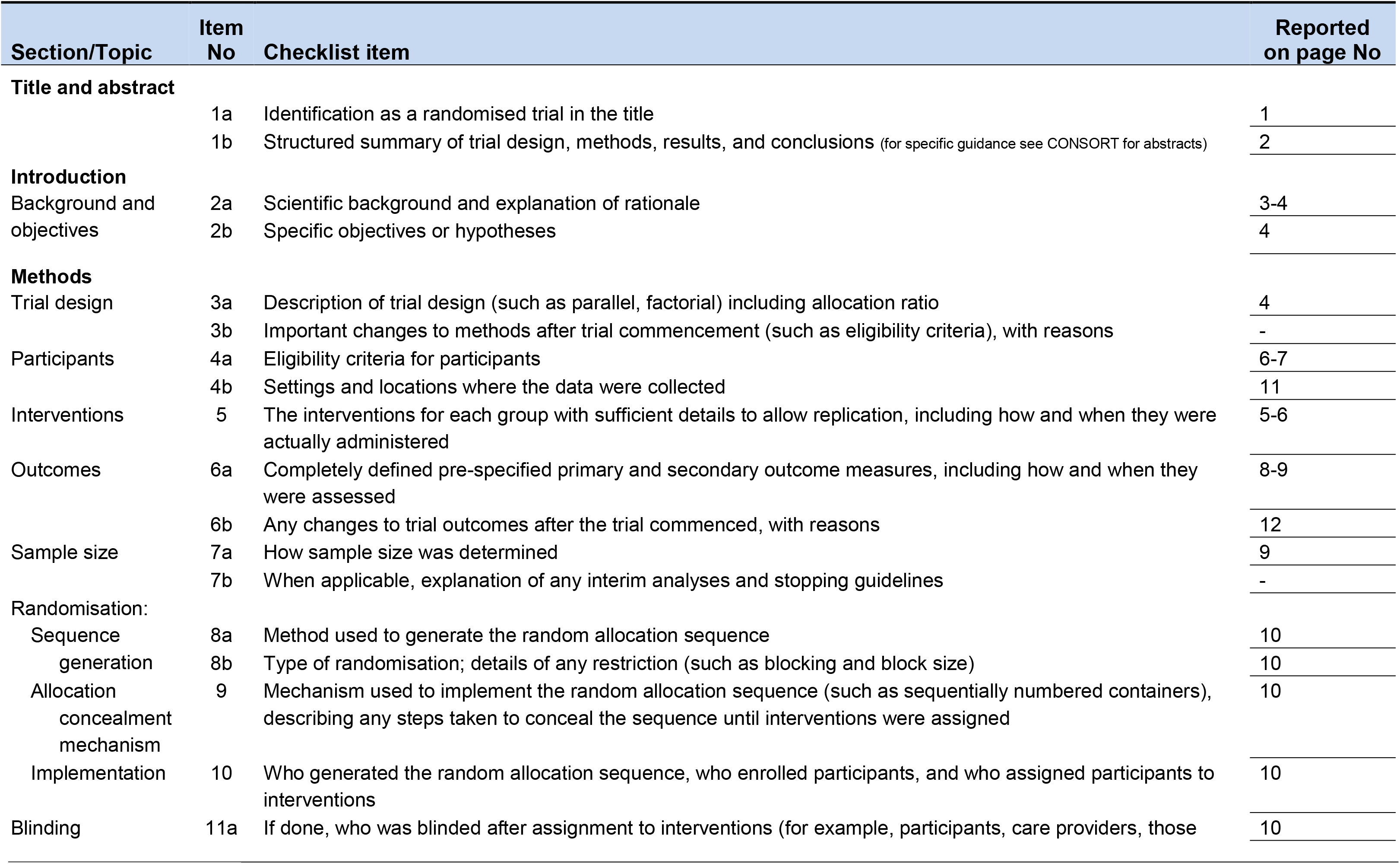
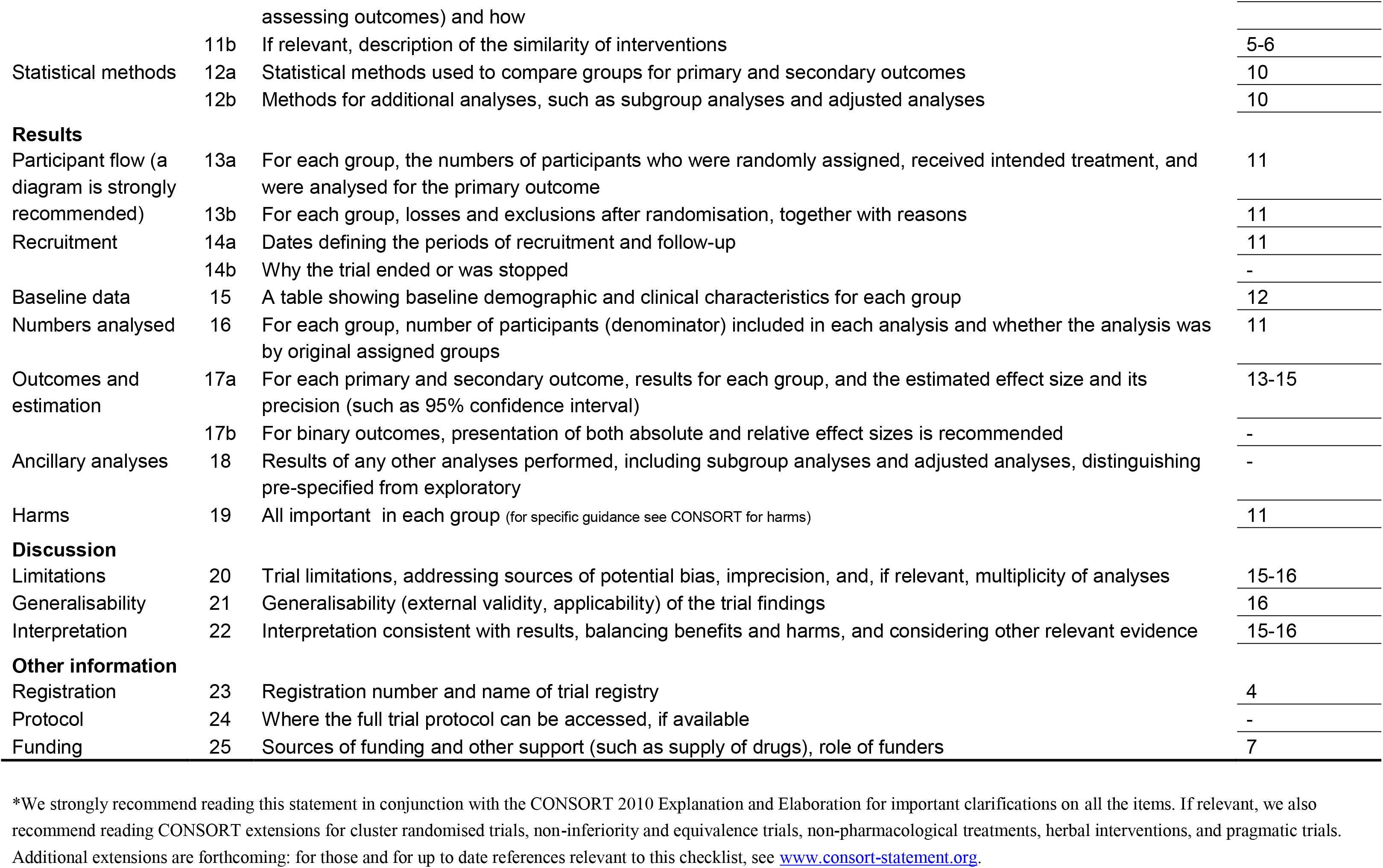
CONSORT Flow Chart CONSORT 2010 checklist of information to include when reporting a randomised trial*.

## Results

### Health Status, Age, Gender

All study patients were healthy. Mean age of the HAP test group patients was 13.4 yrs ± 1.8 SD and 13.4 yrs ± 1.7 SD for the fluoride control group. 52.7% of the HAP test group and 62.2% of the fluoride control group patients were female.

### Blinded change of the primary outcome

A blinded analysis of the ICDAS data at the end of the study revealed, that the overall observed occurrence of ICDAS lesions ≥ code 2 in the study population was 29.3% and therefore considerably lower than the anticipated value (p = 80%) used for the sample size calculation. As the difference between the groups not be regarded clinically relevant had been set in the study protocol to Δ ≤ 20% a clinically meaningful verification of non-inferiority was no longer warranted. Thus, the primary endpoint was changed to the more frequent overall occurrence of ICDAS lesions ≥ code 1 (59.2%). It was decided to keep the original primary endpoint as an additional secondary outcome in the statistical data analysis.

### *Occurrence of ICDAS lesions* ≥ code 1 and ≥ code 2

The occurrence of ICDAS lesions ≥ code 1(revised primary outcome) and ICDAS lesions ≥ code 2 (secondary outcome) is depicted in table 1. In the PP analysis 54.7% of the HAP group patients (n=35) and 60.9% of the fluoride control group patients (n=42) showed the formation of at least one ICDAS lesion ≥ code 1 during the 168 day observation period. In the ITT analysis the corresponding numbers were 56.8% for the patients of the HAP group (n=42) and 61.9% for those of the fluoride control (n=45). In the PP data set the occurence of at least one ICDAS lesion ≥ code 2 was observed in 23.4% of the patients of the HAP group compared to 34.8% of the fluoride controls. In the ITT data set the the corresponding numbers were 25.7% of the HAP group and 32.9% of the fluoride controls. Differences between the groups could not be verified statistically for both analysis sets.

**Table 1.**
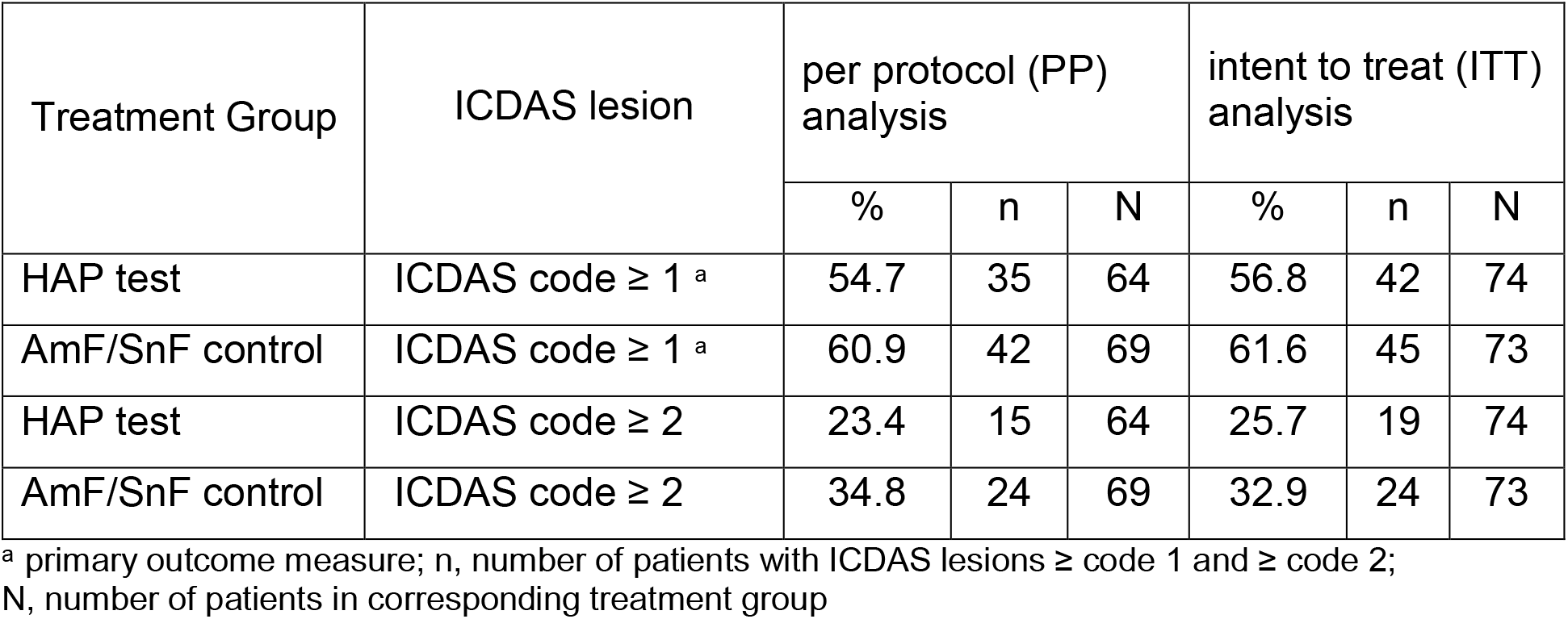
Occurrence of ICDAS lesions ≥ code 1 and ≥ code 2 within the 168 days observation period (ITT and PP analysis)

| Treatment Group | ICDAS lesion | per protocol (PP) analysis |  |  | intent to treat (ITT) analysis |  |  |
| --- | --- | --- | --- | --- | --- | --- | --- |
|  |  | % | n | N | % | n | N |
| HAP test | ICDAS code $\geq 1$ <sup>a</sup> | 54.7 | 35 | 64 | 56.8 | 42 | 74 |
| AmF/SnF control | ICDAS code $\geq 1$ <sup>a</sup> | 60.9 | 42 | 69 | 61.6 | 45 | 73 |
| HAP test | ICDAS code $\geq 2$ | 23.4 | 15 | 64 | 25.7 | 19 | 74 |
| AmF/SnF control | ICDAS code $\geq 2$ | 34.8 | 24 | 69 | 32.9 | 24 | 73 |

<sup>a</sup> primary outcome measure; n, number of patients with ICDAS lesions ≥ code 1 and ≥ code 2; N, number of patients in corresponding treatment group

### Non-inferiority analysis

Table 2 displays the difference between both experimental groups regarding the percentage of study subjects experiencing the new occurrence of at least one ICDAS lesion ≥ code 1 (primary outcome) or at least one ICDAS lesion ≥ code 2 (secondary outcome) including the corresponding one-sided 95% confidence intervals. As the upper limits of the 95% confidence intervals for the primary outcome are well below the given non-inferiority margin of Δ ≤ 20% for both analysis sets (PP: 8%; ITT: 9%) the HAP group has to be considered as non-inferior to the fluoride control.

**Table 2.** Difference between both experimental groups regarding the occurrence of ICDAS lesions ≥ code 1 and ≥ code 2 within the 168 days observation period (95% one-sided confidence intervals)

| Analysis | ICDAS lesion | Proportion in risk difference | Exact lower one-sided 95% confidence limit | Exact upper one-sided 95% confidence limit <sup>b</sup> |
| --- | --- | --- | --- | --- |
| PP analysis | ICDAS code ≥ 1 <sup>a</sup> | -0.062 | -0.203 | 0.083 |
| ITT analysis | ICDAS code ≥ 1 <sup>a</sup> | -0.048 | -0.188 | 0.087 |
| PP analysis | ICDAS code ≥ 2 | -0.114 | -0.255 | 0.030 |
| ITT analysis | ICDAS code ≥ 2 | -0.072 | -0.202 | 0.068 |

<sup>a</sup> primary outcome measure; <sup>b</sup> the upper one-sided 95% confidence limit is markedly lower than the non-inferiority margin of 0.20 ( $\Delta = 20\%$ ) thus inferiority is rejected.

Also regarding the secondary outcome (ICDAS lesion ≥ code 2) the upper limits of the 95% confidence intervals are substantially below the given non-inferiority margin of 20% for both analysis sets (PP: 3%, ITT: 7%), indicating that again the HAP test group has to be considered being non-inferior to the fluoridated control.

### Effect of study site on the primary outcome measure

The effect of study site on the primary outcome measure ΔICDAS score ≥ 1 at week 24 was evaluated by logistic regression analysis. It included the factors study site, treatment group and the interaction between study site and treatment group. Due to small sample sizes, the data for the study sites Dresden, Munich and Frankfurt were pooled (n=40 patients). The results revealed a significantly lower incidence of the primary outcome at week 24 (p<0.001) at the combined smaller centers (Dresden, Munich, Frankfurt) when compared to the study centers in Regensburg (n=72 patients) or Wuerzburg (n=35 patients). However, there was no significant interaction between study site and treatment group, proving that the factor study site did not significantly affect efficacy differences between the treatment groups (for further informations see also Appendix 1).

### Number and severity of ICDAS score increases

The number and severity of ICDAS score increases on the vestibular surfaces of teeth 15-25 over the course of the study are shown in table 3. At week 4 3.2% of the teeth in the HAP group were already affected (ICDAS code 1: 3.0%; ICDAS code 2: 0. 2%) compared to 3.6% of the AmF/SnF control group (ICDAS code 1: 3.1%; ICDAS code 2: 0.5%). These figures steadily increased over time. At week 24 19.6% of the teeth in the HAP group were affected (ICDAS code 1: 14.8%; ICDAS code 2: 4.8%) compared to 21.0% in the AmF/SnF control group (ICDAS code 1: 14.2%; ICDAS code 2: 6.7%; ICDAS code 3: 0.1%).

**Table 3.** Number and severity of ICDAS score increases observed on teeth 15-25 at week 4, week 12 and week 24 (PP data set; n=133)

|  |  | **HAP**<br>**test group** |  | **AmF/SnF**<br>**control group** |  | **Total** |  |
| --- | --- | --- | --- | --- | --- | --- | --- |
| **Visit** | **Δ ICDAS** | **n**<br>**teeth** | **%** | **n**<br>**teeth** | **%** | **n**<br>**teeth** | **%** |
| week 4 | No increase | 620 | 96.9 | 665 | 96.4 | 1285 | 96.6 |
|  | Δ ICDAS code 1 | 19 | 3.0 | 24 | 3.5 | 43 | 3.2 |
|  | Δ ICDAS code 2 | 1 | 0.2 | 1 | 0.1 | 2 | 0.2 |
|  | **Total** | 640 | 100.0 | 690 | 100.0 | 1330 | 100.0 |
| week 12 | No increase | 573 | 89.5 | 611 | 88.6 | 1184 | 89.0 |
|  | Δ ICDAS code 1 | 58 | 9.1 | 59 | 8.6 | 117 | 8.8 |
|  | Δ ICDAS code 2 | 9 | 1.4 | 20 | 2.9 | 29 | 2.2 |
|  | **Total** | 640 | 100.0 | 690 | 100.0 | 1330 | 100.0 |
| week 24 | No increase | 514 | 80.3 | 545 | 79.0 | 1059 | 79.6 |
|  | Δ ICDAS code 1 | 95 | 14.8 | 98 | 14.2 | 193 | 14.5 |
|  | Δ ICDAS code 2 | 31 | 4.8 | 46 | 6.7 | 77 | 5.8 |
|  | Δ ICDAS code 3 | 0 | 0 | 1 | 0.1 | 1 | 0.1 |
|  | **Total** | 640 | 100.0 | 690 | 100.0 | 1330 | 100.0 |

### Plaque Index (PlI), Gingival Index (GI)

The results of the ITT analysis of the PlI and the GI data are shown in Table 4. Mean PlI as well as mean GI scores increased significantly (p < 0.0001) between baseline and day 168 in both groups. Neither at baseline nor at day 168 differences between the experimental groups were statistically significant.

**Table 4.** Plaque Index and Gingival Index scores at baseline, day 28, day 84 and day 168 (ITT analysis)

| Plaque Index (PII) |  |  |  |  |  |  |
| --- | --- | --- | --- | --- | --- | --- |
|  | HAP test group |  |  | AmF/SnF control group |  |  |
| Visit | N | mean<sup>a</sup> | SD | N | mean<sup>a</sup> | SD |
| baseline<sup>b</sup> | 75 | 0.35 | 0.37 | 74 | 0.36 | 0.36 |
| week 4 | 74 | 0.65 | 0.58 | 72 | 0.76 | 0.56 |
| week 12 | 74 | 0.72 | 0.60 | 73 | 0.75 | 0.61 |
| week 24<sup>b</sup> | 74 | 0.85 | 0.66 | 73 | 0.77 | 0.61 |
| Gingival Index (GI) |  |  |  |  |  |  |
|  | HAP test group |  |  | AmF/SnF control group |  |  |
| Visit | N | mean<sup>a</sup> | SD | N | mean<sup>a</sup> | SD |
| baseline<sup>b</sup> | 75 | 0.29 | 0.36 | 74 | 0.37 | 0.41 |
| week 4 | 74 | 0.53 | 0.57 | 73 | 0.58 | 0.54 |
| week 12 | 74 | 0.51 | 0.53 | 73 | 0.66 | 0.55 |
| week 24<sup>b</sup> | 74 | 0.70 | 0.56 | 73 | 0.77 | 0.59 |

<sup>a</sup> significant ( $p < 0.0001$ ) increase of Plaque Index and Gingival Index over time from baseline to day 168 for both treatment groups (Friedman test)
<sup>b</sup> no significant differences between both treatment groups at baseline and at day 168 (two-sided Wilcoxon-Mann-Whitney test)

## Discussion

### Methods

Caries detection and grading in this trial followed the principles of ICDAS-II [23], an internationally established, state of the art caries assessment method, which is particularly suitable and appropriate for the differentiation and grading of incipient enamel caries development, allowing to verify even minor differences in the efficacy of caries-preventive measures. As described before, repeated examiner calibration by an internet-based ICDAS training tools as well as personal examiner calibrations by an experienced ICDAS grading expert (K.H.K.) were integral parts of the study design, to warrant the validity of the clinical recordings. Mean weighted kappa for interrater reliability increased from 0.75 for the first to 0.80 for the final calibration assessment. This was overall in the upper range of kappa reliability scores reported by other controlled clinical trials and indicated “substantial” agreement [25]. It assured that the assessment of the primary study outcome was based on a sound foundation. Furthermore, all examiners were blinded to the dentifrice allocation of the study subjects throughout the course of the study to avoid any possible examiner bias.

### Study population

The assessed study patients, wearing fixed orthodontic appliances, were without doubt caries-active, documented by the considerable increase in enamel caries lesions during the 168 days observation period, which was comparable in its magnitude to observations made by other clinical trials [18, 29]. Due to the inevitable lack of a negative control group for ethical reasons, it is however very difficult to assess the true extent of caries prevention provided by the regular use of both dentifrices to the study participants. It may be argued, that in the chosen setting of caries-active orthodontic patients overly acidic conditions beyond the remineralizing capacities of fluorides and hydroxyapatite particles might have rendered a meaningful non-inferiority analysis impossible. We cannot share, however, this assumption for the following reasons: In a more recent multicenter caries trial by Sonesson et al. (2014), assessing a comparable cohort of 424 adolescent patients age 12-16 subjected to orthodontic therapy with fixed appliances, the regular use of a highly concentrated 5000 ppm fluoride dentifrice was accompanied by a significantly lower incidence of white spot enamel lesions when compared to the regular use of a standard 1450 ppm fluoride control dentifrice [30]. While this suggests, that 1450 ppm may not be the optimal fluoride concentration for a dentifrice to be used in caries-active orthodontic patients, it evidently contradicts the assumption, that in these patients the caries-preventive efficacy of fluoride is completely blocked by overly acidic conditions. On the contrary, Sonesson et al. even speculated, that the reduced increase in enamel caries observed for the use of the 5000 ppm fluoride dentifrice may be attributable to a dose-dependent inhibitory effect of fluorides on *in vivo* lactate production in supragingival bacterial biofilms as discovered by Takahasi and Washio (2012) [31]. The findings by Sonesson et al. are also in line with the conclusions of two meta-reviews assessing the efficacy of topical fluorides in orthodontic patients subjected to therapy with fixed braces, which advocated the additional use of fluoride rinses or varnishes as a complement to regular toothbrushing with fluoride dentifrices [17, 32]. From a clinician’s point of view the caries protection provided by the sole use of both dentifrices evaluated in the present trial may not have been sufficient for a sizeable part of the study participants. This may however not be interpreted as an inherent and complete lack of clinical efficacy. It rather reflects the fact, that the study population deliberately and in accordance with the recommendations of the 2004 International Consensus Workshop on caries clinical trials [33], included many subjects with a very high caries activity. While this is a condition, which clinically may only be partially controllable by the sole use of conventional fluoride toothpaste not exceeding the legal fluoride concentrations limits for cosmetic products, as described before, the inclusion of highly caries-active individuals in a controlled caries prevention trial is an indispendable prerequisite for a meaningful verification of possible differences regarding the caries-protecting capacities of evaluated products or interventions [33].

### Data Analysis

Whether the occurrence frequency of ICDAS code 1 enamel caries lesions used in this study is the most suitable primary endpoint for an non-inferiority caries trial may be subject to discussion. However, the adjunctive analysis of the PP data set regarding frequency and severity of the occurrence of enamel caries lesions during the observation period depicted in table 3 only confirms the identified absence of relevant differences between both experimental groups. It may also have been debatable to keep the original non-inferiority margin of Δ = 20% when switching the primary outcome of the trial despite an overall incidence of the revised primary outcome (ICDAS lesion code 1) of only 60%. The subsequent analysis of the unblinded PP data set however revealed, that the actual difference between both experimental groups was 6.2% in favour of the HAP test dentifrice with an exact upper one-sided 95% confidence limit of 8.3%, i.e. substantially lower than the preset non-inferiority margin of Δ = 20%.

### Secondary Outcomes

The data for the secondary outcomes Plaque Index (PlI) and Gingival Index (GI) furthermore confirmed the findings of preceding studies, reporting a significant increase of gingival inflammation and bacterial plaque mass after the onset of orthodontic therapy with fixed appliances [18, 29]. Differences between both experimental groups regarding the recorded PlI and GI data could not be verified statistically for any of the evaluated time points, which is also in good agreement with the results of a previous trial comparing the plaque- and gingivitis-reducing properties of a fluoride-free HAP test dentifrice and a fluoridated AmF/SnF control in a study cohort of periodontitis patients [11].

### Outlook

While the safety of fluoride-based caries prevention has been firmly established by numerous studies [21], dosage and toxicity aspects have always to be considered. This particularly limits the clinical feasibility of the aforementioned increase in fluoride dosing in high caries-risk infants and children up to an age of 8 due to the asscociated risk for the development of dental fluorosis. Although not verified by clinical studies so far, increasing the dosing or application frequency of HAP toothpaste might also have a beneficial impact on clinical outcome in highly caries active subjects because HAP is a potent buffer, able to neutralize organic acids in a dose-dependent manner, By contrast to fluorides, increasing the applied dosage of HAP particles is virtually free of any toxicity issues even in infants and children, as HAP is the major mineral phase of all hard tissues within the human body [6].

### Conclusions

Regular toothbrushing with a fluoride-free microcrystalline hydroxyapatite-containing dentifrice is a viable method of clinical caries control. The proof of non-inferiority in comparison to the regular use of a conventional fluoride dentifrice verified by this trial in a cohort of caries-active adolescents and adults may however be only a first step. The promising results revealed by the present study need to be corroborated by subsequent clinical investigations in a broader spectrum of study populations and diverging caries activities before general conclusions regarding the benefits and limits of microcrystalline HAP in clinical caries prevention may be possible.

## Acknowledgements

The contract research organisation FGK Clinical Research GmbH, Germany performed the statistical analyses. The statistical analyses, including sample size calculation, were planned by the Society for Biometry and Psychometry, Bielefeld, Germany in cooperation with the principal investigator of the study.

